# *S*-nitrosylation of Aux/IAA protein represses auxin signaling

**DOI:** 10.1101/2022.10.07.511298

**Authors:** Hongwei Jing, Xiaolu Yang, Jian Feng, Jian Zhang, Lucia C. Strader, Jianru Zuo

## Abstract

Auxin plays crucial roles in nearly every aspect of plant growth and development. Auxin signaling activation is mediated through degradation of Auxin/INDOLE-3-ACETIC ACID (Aux/IAA) family. Nitric oxide (NO) regulates diverse cellular bioactivities through *S*-nitrosylation of target protein at specific cysteine residues. NO-auxin interplay has an important role in regulation plant growth. However, little is known about the molecular mechanism of how NO effects Aux/IAA proteins stability. Here we show that NO negatively regulates the IAA17 protein stability to repress auxin signaling. We found that NO directly inhibits IAA17 protein degradation. *S*-nitrosylation of IAA17 at Cys-70 represses the TIR-IAA17 co-receptor interaction to attenuate auxin responsiveness. Our data suggest a model in which *S*-nitrosylation of IAA17 at Cys-70 negatively regulates auxin signaling to effect plant development, providing a mechanism for redox-phytohormones networks.

## INTRODUCTION

The phytohormone auxin plays a pivotal role in various aspects throughout the plant life cycle. The predominant nuclear auxin signaling pathway is mediated by three major components: TRANSPORT INHIBITOR RESPONSE1/AUXIN SIGNALING F-BOX PROTEIN (TIR1/AFB) receptors, AUXIN RESPONSE FACTOR (ARF) transcription factors and Auxin/INDOLE-3-ACETIC ACID (Aux/IAA) repressor proteins^1^. Under low auxin conditions, Aux/IAA proteins directly repress ARF transcriptional activity. When auxin levels increase, auxin promotes complex formation of SCF^TIR1/AFB^ and Aux/IAAs. The SCF^TIR1^-auxin-Aux/IAA co-receptor formation causes polyubiquitylation of Aux/IAA proteins, which are subsequently degraded through the 26S proteasome. Aux/IAA protein degradation relieves ARF protein repression, allowing ARF protein regulation of auxin-responsive genes^2^. Aux/IAA repressor proteins tightly control of auxin transcriptional events. In Arabidopsis, there are 29 Aux/IAA proteins that interact with other auxin signaling components to activate auxin downstream response^3^. Most Aux/IAA proteins contain three homology domains, consisting of an N-terminal domain I (DI) for recruitment co-repressors, followed by degron domain (DII) determines protein stability, and terminating in a C-terminal type I/II Phox and Bem1p (PB1) protein-protein interaction domain^4^.

Post-translational modifications (PTMs) are critical for maintaining the fundamental functions of plant living cells major through regulating protein function, conformation, subcellular localization, interactions, and protein stability^5^. Aux/IAA proteins are modified by multiple types of PTMs, including ubiquitylation, *cis-trans* isomerization, and phosphorylation^3^. Nitric oxide (NO), as a gaseous signaling molecule, plays myriad roles in plant growth and development, including hormone regulation, signaling transduction, biotic and abiotic stress responses^6-8^. The dominant NO bioactivity is in protein *S*-nitrosylation, which covalently addition an NO moiety to a protein reactive cysteine (Cys) thiol (SH) forming an *S*-nitrosothiol (SNO)^7^. *S*-nitrosylation, as a highly conserved redox-based PTMs, plays crucial roles in various physiological and pathological processes by regulation protein activities^9^. In plants, *S*-nitrosylation of target proteins involved in the various phytohormonal network, including synthesis, transport, degradation, and signaling transduction^10,11^. Most recent progresses indicate that NO plays an important role in regulation of phytohormone auxin response, polar transport and also signaling transduction^12-15^. *S*-nitrosylation of auxin receptor TIR1 at Cys140 residue is important for TIR1 function^13^. ARABIDOPSIS SKP1-LIKE (ASK1), an adaptor in the subunits of SKP1–Cullin–F-box (SCF) complex, is S-nitrosylated and involved in SCF^TIR1/AFBs^ assembly^15^. These findings suggest that multiple events of NO bioactivity might be fine-tuning regulation of SCF^TIR1/AFBs^ for auxin signaling transduction. However, the molecular mechanism of how NO regulates Aux/IAA proteins stability remains unknown.

Here we show that NO represses auxin signaling by inhibiting the interaction between receptor TIR1 and Aux/IAA repressor proteins through *S*-nitrosylation of IAA17 at Cys-70 residue, revealing a molecular mechanism behind the complex web of redox-auxin interaction in regulation of plant growth.

## RESULTS

### NO inhibits IAA17 protein degradation

To investigate whether NO effects the Aux/IAA proteins stability, we used the Arabidopsis *HS:AXR3NT-GUS* line, in which a fusion of N-terminal domains I and II of AXR3/IAA17 (AXR3NT) to the β-glucuronidase encoding gene (*GUS*) reporter protein is expressed under control of a heat-shock inducible promoter (*HS*)^16^. After the heat shock treatment at 37 °C for 2 h, the seedlings were then treated with IAA, specific proteasome inhibitor MG132, NO donors *S*-nitrosoglutathione (GSNO) and sodium nitroprusside (SNP). Similar as the previous report, auxin could rapidly induce the degradation of the AXR3 fusion protein (Fig. 1A, Sup_Fig. 1, 2). However, NO donors GSNO and SNP both strongly blocked this IAA-enhanced degradation over the time course, stabilizing the fusion protein to a similar extent as that observed with MG132 (Fig. 1A, Fig_Sup 1, 2), suggesting that NO could stabilize the IAA17 protein which similarly as previous report^14^. To further confirm the results, we quantified the total GUS protein amounts of the *HS:AXR3NT-GUS* line under different treatment by immunoblot analysis. Consistent with the staining results, the NO significantly inhibited the IAA17 protein degradation process (Fig. 1B, C, Sup_Fig. 2). Moreover, GUS enzyme activity of the *HS:AXR3NT-GUS* reporter line were significantly increased after GSNO treatment in a concentration dependent way when comparing with the IAA treatment (Fig. 1D). These results indicate that NO regulates the IAA17 protein degradation.

**Figure 1.**
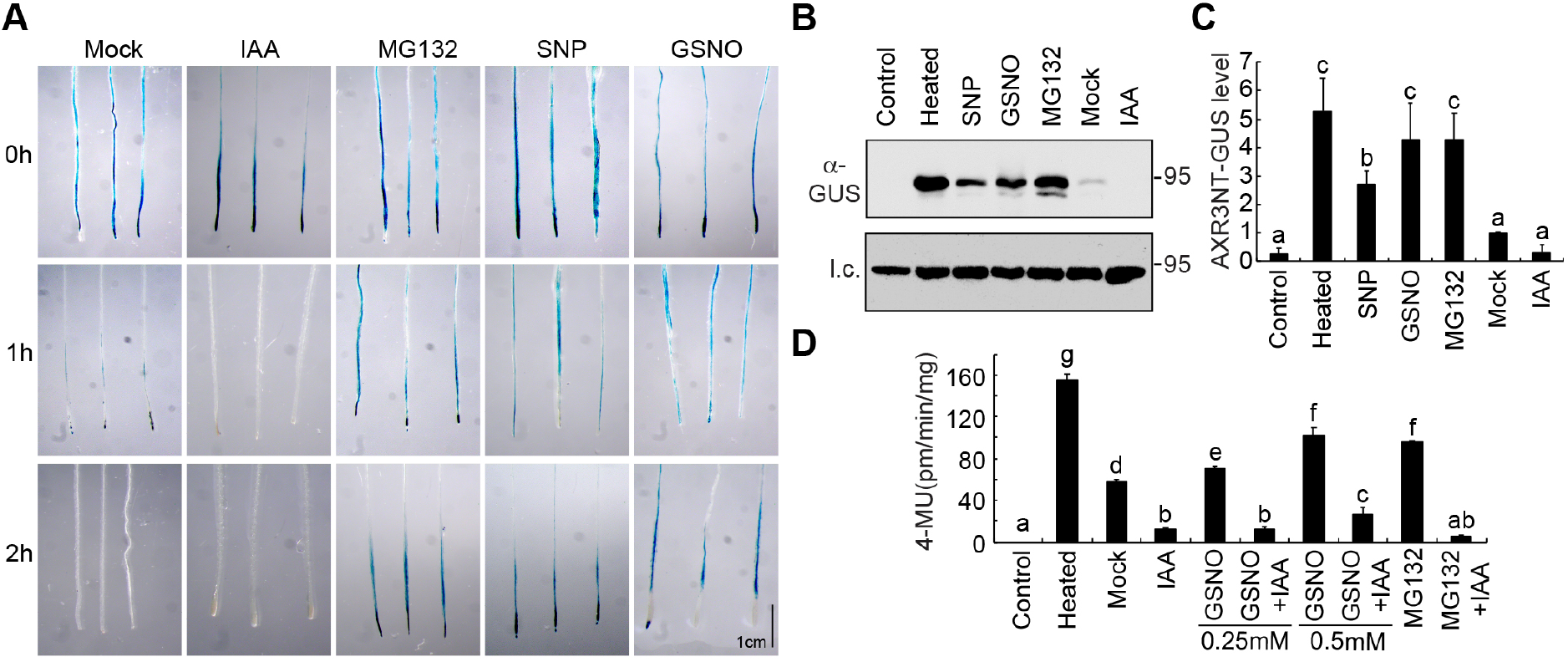
Nitric oxide (NO) regulates IAA17 protein degradation. (**A**) GUS histochemical staining of 7d-old seedlings containing the HS:AXR3NT-GUS/Col-0 reporter. After heat shock at 37 ºC for 2 h, the seedlings were transferred to media in absence (Mock) or presence of 10 μM IAA, 50 μM MG132, 200 μM SNP, and 200 μM GSNO, incubating for 0-2h at 22 ºC before GUS staining. Scale bar = 1cm. See Supplementary Fig. 1 for extended data. (**B, C**) Immunoblot analysis and quantification of GUS levels from 7d-old HS:AXR3NT-GUS/Col-0 seedlings treated with IAA, MG132, SNP, and GSNO with indicated concentrations after the end of the heat shocked at 37 ºC for 2 h. Anti-HSP82 used for loading control (l.c.). (**D**) GUS activity analysis of 7d-old HS:AXR3NT-GUS/Col-0 seedlings treated with IAA, GSNO, MG132 with indicated concentrations after the end of the heat shock period. GUS activity was measured by fluorometric assays. Data are mean ± SD of three independent experiments. Different letters indicate individual groups for multiple comparisons with significant differences (one-way ANOVA, Duncan, p < 0.05).

**Figure 2.**
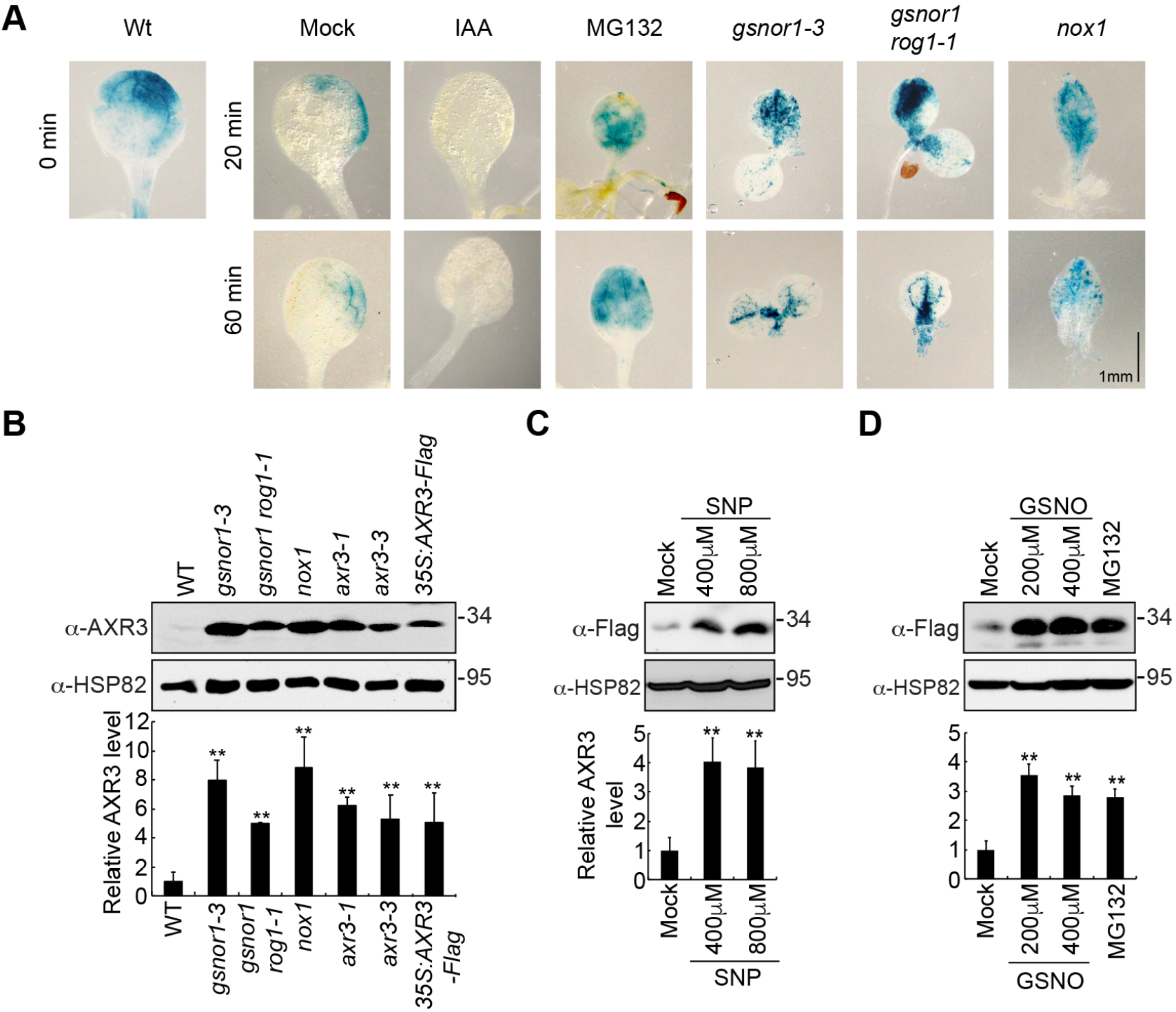
Nitric oxide (NO) regulates IAA17 protein accumulation. (**A**) GUS staining of 7d-old seedlings containing the HS:AXR3NT-GUS reporter in Col-0 or NO-overproducing mutants (*gsnor1-3, gsnor1 rog1-1*, and *nox1*) background after the end of heat-shocked at 37 ºC for 2 h. Scale bar = 1mm. (**B**) Immunoblot analysis (top) and quantification (bottom) of IAA17 protein levels in NO-overproducing mutants *gsnor1-3, gsnor1 rog1-1*, and *nox1*. IAA17 gain of function mutant *axr3-1, axr3-3* and overexpression transgenic plants *35S:AXR3-Flag* were used as positive control. (**C**) Immunoblot analysis (top) and quantification (bottom) of IAA17 protein levels after Nitric oxide (NO) donor SNP treatment. (**D**) Immunoblot analysis (top) and quantification (bottom) of IAA17 protein levels after Nitric oxide (NO) donor GSNO treatment. MG132 was used as positive control. Data are mean ± SD of three independent experiments. The statistical significance was determined by a two-sided Student’s *t*-test (Paired two sample for means). \*\**P* < 0.01 when compared to the mock.

We next examined the IAA17 protein accumulation in the Arabidopsis NO-overproducing mutants, *nox1* (NO overproducer1), *gsnor1-3* (GSNO reductase1), and *gsnor1 rog1-1* (*repressor of gsnor1*). After crossing these NO-overproducing mutants to the *HS:AXR3NT-GUS* background and then heat shock treated at 37 °C for 2 h, all the mutants displayed higher accumulation of AXR3NT fusion protein when comparing with the Mock over the time course (Fig. 2A), further supporting our above data (Fig. 1A, Fig_Sup 1, 2). To further investigate the IAA17 protein accumulation in these NO-overproducing mutants, IAA17 antibodies were generated from the mice using heterologously-expressed His-IAA17 (Sup_Fig. 3). We found that the accumulation of IAA17 were increased in *nox1, gsnor1-3*, and *gsnor1 rog1-1*, displaying a similar way as that observed in the IAA17 gain of function mutant (*axr3-1* and *axr3-3*) and overexpression lines (*35S:AXR3-Flag*) (Fig. 2B). A similar phenotype was also observed in the overexpression lines treated with the NO donors GSNO and SNP (Fig. 2C, 2D). These results demonstrate that accumulation of IAA17 was increased in the NO-overproducing mutants, suggesting NO negatively regulates the IAA17 protein stability.

### IAA17 is *S*-nitrosylated at Cys-15 and Cys-70

Given that NO inhibits the IAA17 protein degradation, a possible mechanism is via the *S*-nitrosylation of cysteine residues in the target proteins. To determine if IAA17 could be *S*-nitrosylated, a biotin-switch method was used to examine the heterologously-expressed IAA17 protein. In this method, the nitrosothiol groups are replaced by a more stable biotin moiety via chemical reduction by ascorbate, and then identified using immunoblot analysis. We found that His- or GST^4CS^-tagged IAA17 recombinant proteins was specifically *S*-nitrosylated by GSNO, but not by glutathione (GSH) (Fig. 3A, 3B). Moreover, S-nitrosylation of the transgenic IAA17-Flag protein was specifically detected *in planta* under normal growth conditions by an *in vivo* biotin-switch method (Fig. 3C).

**Figure 3.**
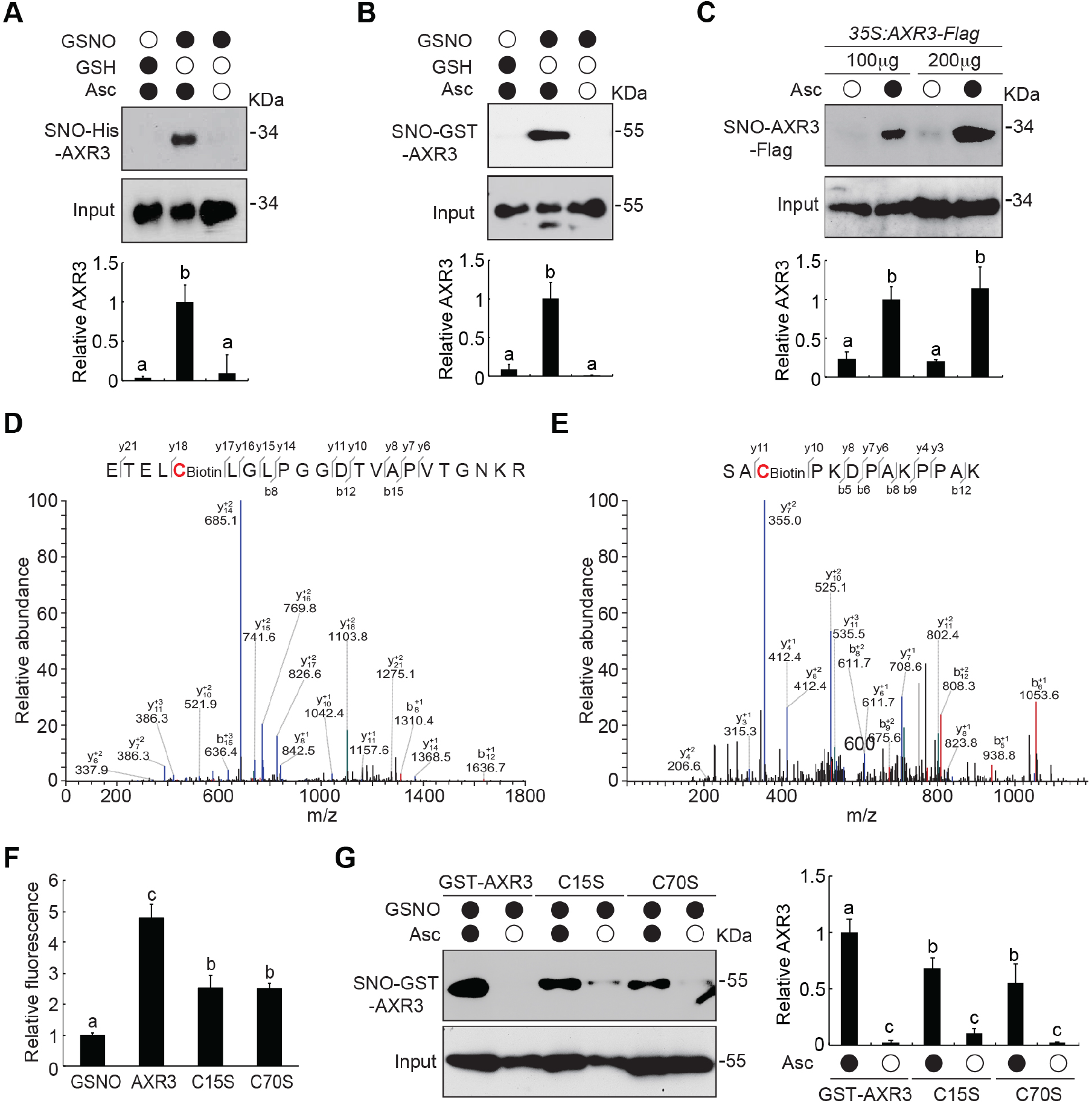
IAA17 is *S*-nitrosylated at Cys-15 and Cys-70. **(A)** Immunoblot analysis (top) and quantification (bottom) of S-nitrosylation of recombinant His-tagged IAA17 (SNO) *in vitro*. Treatment with GSH in absence or presence of ascorbate sodium (Asc) as negative controls. **(B)** Analysis of the S-nitrosylation of recombinant GST^4CS^-tagged IAA17 (SNO) *in vitro*. Quantitative analysis of the data is shown in bottom. Treatment with GSH in absence or presence of ascorbate sodium (Asc) as negative controls. (**C**) *In vivo* analysis of S-nitrosylation IAA17-Flag protein (SNO) in extracts prepared from *35S:AXR3-Flag* seedlings. Quantitative analysis of the data is shown in button. (**D, E**) Analysis of the S-nitrosylation sites of IAA17 by Mass spectrometry (MS/MS) methods from a biotin-charged IAA17 peptide. The b- and y-type product ions are indicated, thereby identifying Cys-15 and Cys-70 as the *S*-nitrosylated residues. (**F**) DAN analysis of *S*-nitrosylated residues in GSNO tripeptide and His-IAA17 recombinant protein. (**G**) Analysis of *S*-nitrosylated GST^4CS^-IAA17, GST^4CS^-IAA17^C15S^, and GST^4CS^-IAA17^C70S^ recombinant proteins. Quantitative analysis of the data is shown in the right. Data are mean ± SD of three independent experiments. Different letters indicate individual groups for multiple comparisons with significant differences (one-way ANOVA, Duncan, *p* < 0.05).

To further identify the *S*-nitrosylated cysteine residues of IAA17, a liquid chromatography-tandem mass spectrometry experiment was performed to analysis the tryptic fragments derived from GSNO-treated IAA17 recombinant proteins. We found that the Cys-15 and Cys-70 as the *S*-nitrosylated residues in IAA17 (Fig. 3D, 3E). In addition, an experiment that measures the conversion ratio of 2, 3-diaminonaphthalene (DAN) into fluorescent 2, 3-naphthltriazole (NAT) catalyzed by NO released from the thiol group, was performed to determine the number of the S-nitrosylated cysteine residues in IAA17. Comparing with the GSNO treatment which contains a single S-nitrosylated Cys residue, IAA17 recombinant protein displayed around 4-flod higher DAN-NAT conversion ratio, whereas both IAA17^C15S^ and IAA17^C70S^ mutant proteins significantly reduced the number of S-nitrosylated Cys residue, suggesting there are more than two *S*-nitrosylated Cys residue existing in IAA17 protein (Fig. 3F). Consistently, IAA17^C15S^ and IAA17^C70S^ mutation protein showed a significantly reduction of S-nitrosylation level comparing with the wild-type IAA17 protein (Fig. 3G). Taken together, these results indicate that Cys-15 and Cys-70 of IAA17 are *S*-nitrosylated sites.

### *S*-nitrosylation negatively regulates the degradation of IAA17

Data presented above indicate that NO regulates the IAA17 protein stability. Since Aux/IAA proteins are degraded by SCF^TIR1^-Aux/IAA receptor formation through the 26S proteasome, we next assessed the possible regulatory effects of NO on the TIR1-Aux/IAA interaction by a co-immunoprecipitation (Co-IP) assay. Considering the TIR1 and ASK1 could be S-nitrosyalted^13,15^, we first incubated the His-IAA17 recombinant protein with GSNO for 2h, then removed the excess GSNO. TIR1-Myc protein purified by immunoprecipitation from transgenic plants was incubated with or without GSNO-treated His-IAA17 recombinant proteins, and the TIR1-IAA17 interaction was examined by a Co-IP experiment. The interaction between TIR1-Myc and His-IAA17 was significantly enhanced by auxin, similarly as the previously observed in Arabidopsis (Fig. 4A). However, when incubated with the GSNO-treated His-IAA17, the interaction between TIR1-Myc and His-IAA17 was dramatically decreased ∼70% when comparing with the auxin induced, similarly as in the absence auxin treatment (Fig. 4A). These results implied that the NO negatively regulated the interaction of TIR1-IAA17 co-receptor in auxin signaling.

**Figure 4.**
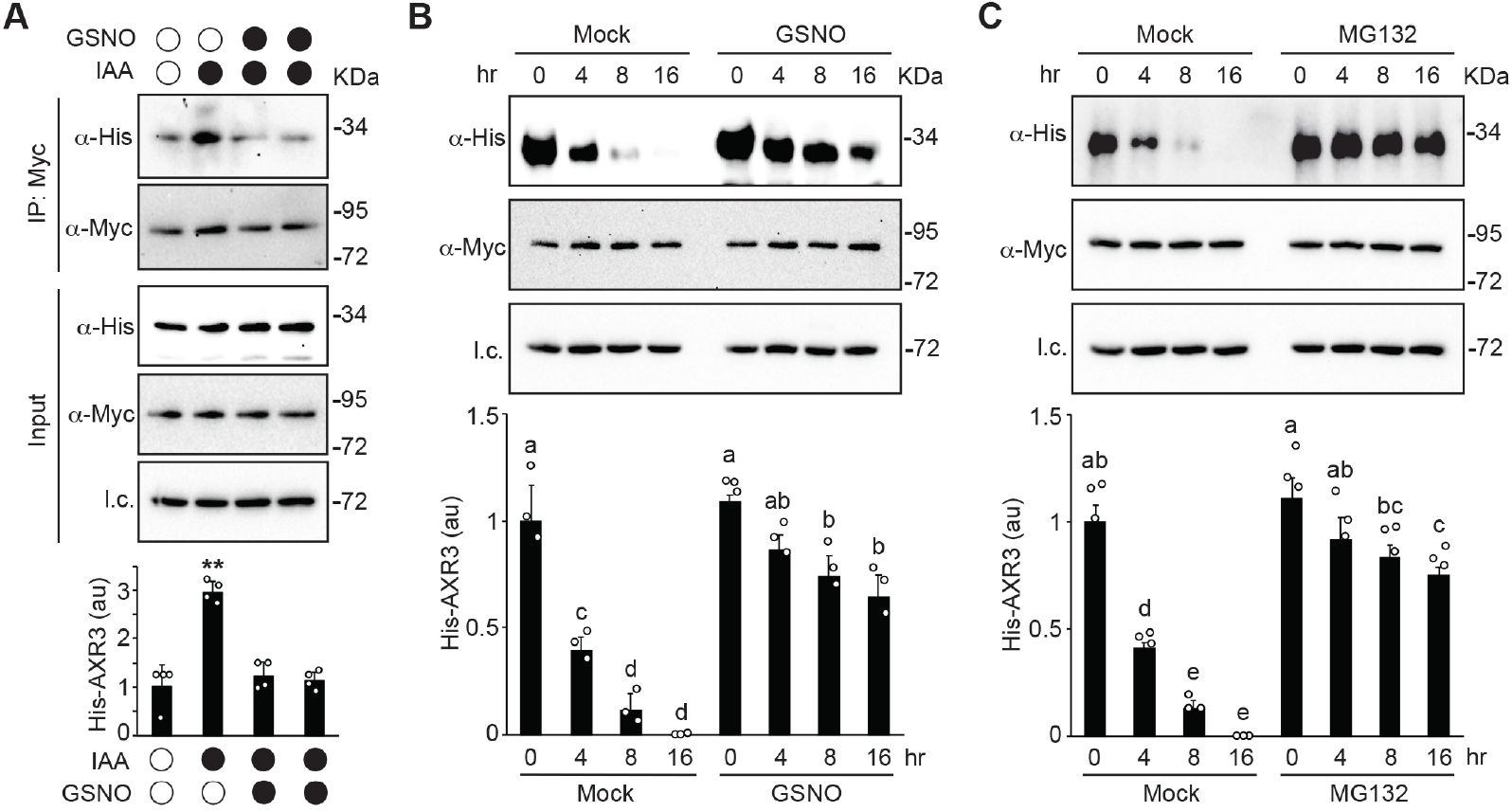
*S*-nitrosylation negatively regulates IAA17 protein degradation. **(A)** Nitric oxide (NO) donor GSNO negatively regulates the auxin-induced TIR1-IAA17 interaction. Firstly, His-IAA17 recombinant protein was incubated with GSNO for 2h, and then removed the excess GSNO by washing. Secondly, the total protein extracts from TIR1-Myc transgenic plants were purified by immunoprecipitation, and was incubated with or without GSNO-treated His-IAA17 recombinant proteins. Then, the immunoprecipitates were analyzed by immunoblotting using antibodies as indicated. Quantitative analysis of the relative level of His-IAA17 is presented below the blots. Data are mean ± SD of three independent experiments. The statistical significance was determined by a two-sided Student’s *t*-test (Paired two sample for means). \*\**P* < 0.01 when compared to the mock. (**B**) *In vitro* His-IAA17 recombinant protein degradation. His-IAA17 recombinant protein was incubated with GSNO for 2h, and then removed the excess GSNO by washing. Then, the total protein extracts from *TIR1-Myc* transgenic plants were incubated with GSNO-treated His-IAA17 recombinant proteins for the indicated times. Immunoblot analysis (top) and quantification (bottom) of His-IAA17 using the indicated antibodies. Anti-HSC70 used for loading control (l.c.). (**C**) Analysis of His-IAA17 recombinant protein degradation *in vitro*. The total protein extracts from *TIR1-Myc* transgenic plants were incubated with His-IAA17 recombinant proteins after adding 50 μM MG132 for indicated times. Immunoblot analysis (top) and quantification (bottom) of His-IAA17 using the indicated antibodies. Anti-HSC70 used for loading control (l.c.). Data are mean ± SD of three independent experiments. Different letters indicate individual groups for multiple comparisons with significant differences (one-way ANOVA, Duncan, *p* < 0.05).

To explore the functional effects of NO on the SCF^TIR1^-Aux/IAA co-receptor formation, we next performed an *in vitro* degradation assay to examine the stability of IAA17. When incubated the transgenic plant lysate of *TIR1-Myc* with the His-IAA17 recombinant proteins, NO donors GSNO remarkably reduced the degradation of His-IAA17 comparing with the mock over the time course, displaying a similar response to MG132, a specific inhibitor of the proteasomal degradation pathway (Fig. 4B, 4C). These results demonstrate that *S*-nitrosylation negatively regulates the TIR1-IAA17 co-receptor formation, thereby reduced the degradation of IAA17 transcriptional repressors through proteasomal pathway.

### *S*-nitrosylation of IAA17 at Cys-70 attenuated auxin responsiveness

Data present above indicate that IAA17 is S-nitrosylated at Cys-15 and Cys-70. To explore the functional role of IAA17 at Cys-15 and Cys-70 on the auxin responsiveness, the transgenic plants *IAA17:IAA17-Flag, IAA17:IAA17*^*C15S*^*-Flag, IAA17:IAA17*^*C15W*^*-Flag, IAA17:IAA17*^*C70S*^*-Flag*, and *IAA17:IAA17*^*C70W*^*-Flag*, carrying wild-type or point mutation *IAA17* cDNA fragments fused to a Flag tag under the control of the native *IAA17* promoter, were generated in the *iaa17* (Salk_011820) T-DNA insertion mutant background. Serine (S), structurally similar to cysteine, has been shown a non-nitrosylation, but tryptophan (W) has been shown to mimic an S-nitrosylated cysteine^17,18^. All the transgenic plants were screened by qRT-PCR experiment, which displaying similar expression levels compared with that of wild type (Sup_Fig. 4).

Analysis the root elongation showed that only *IAA17:IAA17*^*C70W*^*-Flag* transgenic plants displayed resistance to the inhibitory effects of the synthetic auxin 2,4-D (Fig. 5A, 5B, Sup_Fig. 5). In addition, only *IAA17:IAA17*^*C70W*^*-Flag* transgenic plants displayed reduced the lateral root density under 2,4-D treatment (Fig. 5C). Moreover, the *35S:IAA17*^*C70W*^*-Flag* transgenic plants showed significant growth and development phenotype comparing with the wild-type (Sup_Fig. 6). Although there is no significant difference of Cys-15 mutation transgenic plants comparing with the wild type, whether Cys-15 residue involved in the other stress response or environmental changes need to be explored in the future. Taken together, our data suggest that *S*-nitrosylation of IAA17 at Cys-70 plays a critical role in regulation of the auxin response.

**Figure 5.**
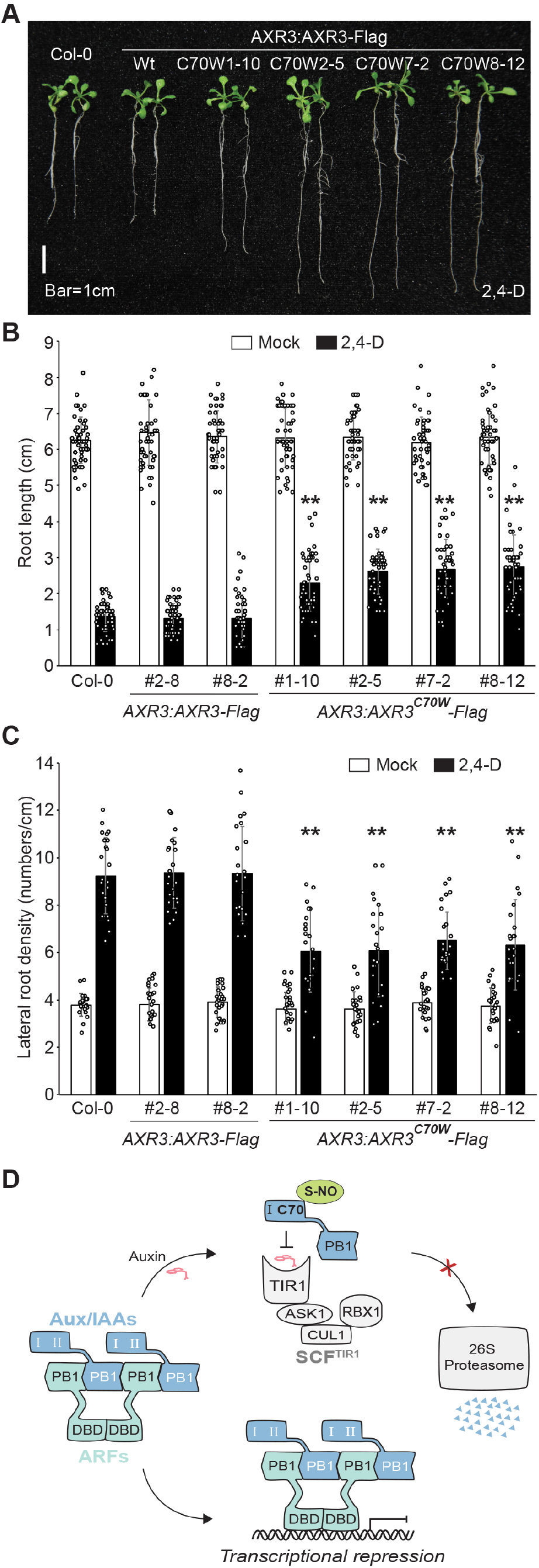
*S*-nitrosylation of IAA17 at Cys-70 residue attenuated auxin responsiveness. **(A)** Photograph of 6d-old Col-0, *AXR3:AXR3-Flag*, and *AXR3:AXR3*^*C70W*^*-Flag* seedlings grown on media supplemented with 10 nM 2,4-D after growing 5 days on 1/2 MS media with sucrose. Scale bar = 1 cm. (**B, C**) Mean primary root lengths and lateral root density of 6d-old Col-0, *AXR3:AXR3-Flag*, and *AXR3:AXR3*^*C70W*^*-Flag* seedlings grown on media supplemented with mock (EtOH) or 10 nM 2,4-D after growing 5 days on 1/2 MS media with sucrose. n = 50 and n=20 biologically independent seedlings were examined in root lengths and lateral root density, respectively. Data are mean ± SD from three independent experiments and gray dots represent the individual values. The statistical significance was determined by a two-sided Student’s *t*-test (Paired two sample for means). \*\**P* < 0.01 when compared to Col-0. (**D**) A proposed model for *S*-nitrosylation of IAA17 at Cys-70 negatively regulates auxin signaling. *S*-nitrosylation of IAA17 at Cys-70 increase the IAA17 protein accumulation by repressing the TIR1-IAA17 co-receptor interaction, thereby attenuating auxin responsiveness to affect plant growth and development.

Overall, our data support a model (Fig. 5D) in which *S*-nitrosylation of IAA17 at Cys-70 modulates IAA17 protein accumulation by repressing the TIR1-IAA17 co-receptor interaction, as a result to regulate auxin responsiveness to affect plant growth and development.

## DISCUSSION

Post-translational modification by NO takes place majorly through three modification, *S*-nitrosylation (the formation of a nitrosothiol group on cysteine residues), metal nitrosylation (interaction of NO with metalloproteins), and tyrosine nitration (covalently addition of NO to the tyrosine residues)^10^. According on current progress, *S*-nitrosylation is considered to play crucial roles in regulation protein activities, that especially involved in phytohormonal network at various levels. NO-auxin interplay that involved in a myriad of morphological and physiological processes, has been shown more complexity and diversity during plant growth and development ^19-21^. More recent studies show that NO treatment reduce the root meristem size and inhibit the root growth by reducing PIN-dependent auxin transport in Arabidopsis^12,22^, suggesting NO impairs auxin polar transport pathway. However, other studies demonstrate that NO directly targets auxin receptor TIR1 for S-nitrosyaltion its Cys-140 residue to enhance the interaction between TIR1 and Aux/IAA repressors^13^, indicating NO could enhance the auxin signaling transduction. Moreover, NO enhances ASK1 by *S*-nitrosyaltion at Cys-37 and Cys-118 residues binding to the subunits of SCF complexes impacting on auxin signaling^15^. In this study, we found that NO directly inhibit the *HS:AXR3NT-GUS* protein degradation, similar as previous report^14^. Genetics and biochemistry evidence demonstrate that NO regulates the proteasome degradation of IAA17 by S-nitrosyaltion at Cys-70 residue, which represses the interaction of TIR1 and Aux/IAA protein to attenuate auxin responsiveness.

The Arabidopsis IAA17 contains five cysteine residues, two of them (Cys15 and Cys70) were identified as *S*-nitrosylation sites by our LC-MS/MS and mutagenesis studies, and DAN-NAT experiments suggested that there are more than two *S*-nitrosylated Cys residue existing in IAA17. In addition, Cys70 is not conserved in other Aux/IAA proteins in Arabidopsis, raising the possibility that *S*-nitrosylation at Cys70 specifically regulates the IAA17 protein stability. More studies relating other cysteine roles in IAA17 and whether other Aux/IAA protein modified by *S*-nitrosylation need to be explored in the further. Although the specificity of *S*-nitrosylation IAA17 at Cys70 inhibit the interaction between TIR1 and IAA17, NO plays a negative role in the regulation of the other phytohormone signaling pathways, suggesting the possibility of existing a general similar mechanism as observed in here that NO-repressed auxin signaling. For instance, NO negatively regulates gibberellin signaling by promoting the accumulation of the gibberellin signaling repressor DELLA in Arabidopsis^23^. Recent studies report that NO induces the *S*-nitrosylation of DELLA protein REPRESSOR-OF-*ga1-3* (RGA) at Cys-374 to inhibit the RGA-SLEEPY1 (SLY1) interaction, thereby negatively regulates gibberellin signaling to coordinate plant growth and salt stress tolerance^24^. In the cytokinin signaling, *S*-nitrosylation of the histidine phosphotransfer protein (AHP1) at Cys-115 to repress its phosphorylation and subsequent transferring of the phosphoryl group, thereby resulting in a compromised cytokinin response^17^. Further, NO chemically reacts with the cytokinin (zeatin), which directly causes the reduction of cytokinin species, further supporting a negative regulatory role of NO in cytokinin pathway^25^. In addition, regarding the environmental stress, NO inhibits the sucrose non-fermenting 1 (SNF1)-related protein kinase (SnRK2.6)/open stomata 1 (OST1) by *S*-nitrosylation at Cys-137 to negatively regulates the abscisic acid (ABA) signaling in the stomata guard cells^26^. Moreover, *S*-nitrosylation of JASMONATE ZIM DOMAIN (JAZ1) repressor proteins at Cys-229 blocks its interaction with the Jasmonic acid (JA) signaling receptor CORONATINE-INSENSITIVE 1 (COI1), thereby resulting in the suppression of JA signaling^27^. Taken together, these observations raise the possibility that NO plays critical roles in negatively regulation of phytohormones signaling pathways depending on the specific developmental stages or different environmental responses. Thus, it seems likely that the NO-modulated signaling transduction of phytohormones appears to be a conserved regulatory mechanism, coordinating plant growth and development in response to dynamically environmental cues.

In summary, our biochemical and genetic evidence suggest that NO negatively regulates the auxin signaling through *S*-nitrosyaltion of IAA17 at Cys-70, thereby repressing its interaction with the auxin receptor TIR1 to attenuate auxin response, providing a general mechanism behind the complex web of redox-phytohormones networks under the various physiological and environmental conditions.

## Supporting information

Supplemental data will be used for the link to the file on the manuscript

## ACKNOWLEDGMENTS

We thank Dr Mark Estelle for the *HS:AXR3NT-GUS* and *TIR1-Myc* seeds; Dr Gary Loake for *gsnor1-3* seeds; Dr Yikun He for *nox1* seeds; the Arabidopsis Biological Resource Center (ABRC) for *axr3-1, axr3-3*, SALK_011820, SALK_065697 seeds; the members of Zuo lab and Strader lab for the helpful discussion. This work was supported by grants from National Natural Science Foundation of China (31830017).

## AUTHOR CONTRIBUTIONS

H.J., X.Y., J.F., and J.Zuo designed the experiments. H.J. and X.Y performed the experiments, assisted by J. Z.. H.J. and X.Y. wrote the manuscript, assisted by L.C.S. and J.Zuo.

## METHODS

### Plant material and phenotypic assays

All *Arabidopsis thaliana* lines were in the Columbia (Col-0) background in all experiments. The *HS:AXR3NT-GUS* ^16^ and *TIR1-Myc* ^28^ transgenic lines were provided by Mark Estelle. The *gsnor1-3* and *nox1* were provided by Gary Loake and Yikun He, separately. The *gsnor1 rog1-1* was reported in our previous work^29^. The *IAA17* gain of function mutant (*axr3-1* and *axr3-3*) and T-DNA insertion mutant (Salk_011820 and Salk_065697) were obtained from ABRC.

For phenotypic assays, seeds were surface sterilized for 15 min with 20% (v/v) bleach with 0.01% (v/v) Triton X-100, then rinsed four times with sterile water. Sterilized seeds were stratified for 2 d at 4ºC to promote uniform germination. After stratification, seeds were plated on 1/2 MS media solidified with 1% (w/v) sucrose at 22 ºC under continuous illumination. To examine root elongation in Col-0 and site mutation transgenic plants, firstly, the seedlings were grown on the 1/2 MS media solidified with 1% (w/v) sucrose for 5 d of vertical growth at 22 ºC under continuous illumination, then the seedlings showing similar root length, were transferred to the media supplemented with mock (EtOH) or the indicated concentration of 2,4-D. The root lengths and lateral root density were measured after 6 d of vertical growth at 22 ºC under continuous illumination.

### Vector construction and plant transformation

To create the pHis-IAA17 and pGST^4CS^-IAA17 expression vectors, the coding sequences of IAA17 was PCR amplified from Arabidopsis cDNA and the stop codons was eliminated during PCR. The PCR fragments with appropriate restriction sites was cloned into the *Bam*HI and *Sal*I sites of pET28a (Novagen) and pGST^4CS^. The pGST^4CS^ was generated from pGST4T-1 (Amersham Biosciences), which showing the *S*-nitrosylation of GST^4CS^ is eliminated^17^. The pGST^4CS^-IAA17^C15S^ and pGST^4CS^-IAA17^C70S^ were generated as a same way.

To construct *35S::IAA17-Flag* vector, the coding sequences of IAA17 was PCR amplified from Arabidopsis cDNA and the stop codons was eliminated during PCR. The PCR fragments with appropriate restriction sites were digested with *Sma*I and cloned into the *Sma*I site of a pSK-Flag vector to generate *IAA17-Flag* fusion genes. The *IAA17-Flag* fusion genes were cloned into the *Kpn*HI and *Sal*I sites of pWM101 under the control of 35S promoter. The *35S::IAA17*^*C70W*^*-Flag* vector was generated as a same way.

To generate the *IAA17::IAA17-Flag* vector, the genomic fragments of *IAA17* that include the putative promoter sequences and the coding regions were amplified by PCR using wild-type Arabidopsis genomic DNA. In these PCR fragments, the stop codons were eliminated and the appropriate restriction sites were introduced during PCR. The PCR fragments with appropriate restriction sites were digested with *Sma*I and cloned into the *Sma*I site of a pSK-Flag vector to generate *IAA17::IAA17-Flag* fusion genes. These fusion genes were released by *Sal*I and *Bam*HI digestion and cloned into the same restriction sites of the vector pBI121, which modified by replacing the 35S promoter with the multiple cloning site (MCS) to remove the overexpression promoter. The *IAA17::IAA17*^*C15S*^*-Flag, IAA17::IAA17*^*C15W*^*-Flag, IAA17::IAA17*^*C70S*^*-Flag, IAA17::IAA17*^*C70W*^*-Flag* vectors were generated as the same way.

All the site-directed mutagenesis indicated above were performed using the Easy Mutagenesis System (TransGen Biotech, Beijing), according the manufacturer’s instructions. All PCR-amplified fragments in the constructs described above were verified by extensive restriction digests and further confirmed by DNA sequencing analysis. All the binary expression vectors were introduced into *Agrobacterium tumefaciens* strain GV3101. Transgenic plants were generated via *Agrobacterium*-mediated transformation^30^. All primers used for plasmid construction are listed in Supplementary Table 1.

### Heat induction and GUS activity assay

*HS::AXR3NT-GUS* plants were crossed into the *gsnor1-3, nox1* and *gsnor1 rog1-1* mutant background. Seven-d-old seedlings were submerged in liquid 1/2 MS media and heat shocked for 2 h at 37 ºC. After the heat shock, seedlings were transferred to media in absence (Mock) or presence of 10 μM IAA, 50 μM MG132, 200 μM SNP, and 200 μM GSNO, incubating for 0-2h at 22 ºC before GUS staining. For the GUS enzyme activity assay^16^, seedlings were collected after 1 h treatment and stored in liquid nitrogen until protein extraction. The fluorometric assays were performed by incubating sample extracts in 2 mM MUG (4-methylumbelliferyl-β-D-glucoronide), 50 mM KPO4 (pH 7.0), 0.1% SDS, 0.1% Triton X-100, 10 mM β-mercaptoethanol and 10 mM EDTA for 16 h followed by analysis with a Microplate Reader. Extracts were prepared from ten seedlings and data were normalized against total protein levels.

### Antibody preparation and immunoblot analysis

To prepare anti-IAA17 antibodies, the full-length cDNA fragments of IAA17 were used to produce recombinant proteins tagged with 6 X His. The purified His-IAA17 recombinant proteins were used to immunize mice and rabbits, respectively. Immunoblot analysis was performed as described previously^31,32^. Total plant proteins were prepared by grinding seedlings in liquid nitrogen and then extracted in lysis buffer (50 mM Tris-HCl, pH 8.0, 150 mM NaCl, 1% (v/v) Nonidet P-40, 0.5% (w/v) sodium deoxycholate, 0.1% (w/v) SDS and 1 mM phenylmethylsulfonyl fluoride (PMSF), and 1% (v/v) protease inhibitors cocktail (Sigma-Aldrich, P9599). After centrifugation twice at 13,000 g for 10 min at 4 °C, the supernatant was collected and then subjected to SDS-polyacrylamide gel electrophoresis. After the proteins were electrically transferred onto a PVDF membrane, and then detected with a primary antibody of indicated (usually at 1: 5,000 dilution). The blot was incubated with a secondary antibody (HRP-conjugated goat anti-rabbit IgG or HRP-conjugated goat anti-mouse IgG; Beijing Dingguo Changsheng Biotechnology) at 1: 50,000 dilution. The signal was detected using a SuperSignal Western Femto Maximun Sensitivity Substrate kit (Thermo Scientific, Cat no.: 34096) according to the manufacturer’s instructions. Arabidopsis HSP82 (Beijing Protein Innovation, Cat # K2010) was used as a loading control. In most immunoblotting experiments, the blot was first probed with an antibody specific to a target protein and then stripped, followed by re-probing the blot with anti-HSP82 antibody. Occasionally, two technical replicates (gels) were simultaneously performed and then analyzed with antibodies against the specific targets and HSP82, respectively. The target or loading control bands were quantified using FIJI (ImageJ) and the mean values of 3–5 independent experiments were presented with statistical analysis (one-way ANOVA or Student’s *t*-test) of significant differences when applicable.

### *In vitro S*-nitrosylation Assay

Analysis of the *in vitro S*-nitrosylation modification was performed by a biotin-switch method as described previously^29^. In brief, approximately 5 μg of recombinant protein was incubated with GSH or GSNO (200 μM) in the dark condition at room temperature for 30 min. The sample of nearly 100 μL reaction volume was precipitated with three volumes of cold acetone, and then washed with acetone for three times and resuspended in 300 μL blocking buffer (250 mM Hepes, pH 7.7, 4 mM EDTA, 1 mM neocuproine, 2.5% (w/v) SDS and 200 mM S-methylmethane thiosulfonate). The reaction was carried out at 50 ºC for 1h with frequently vortex. Then the sample were precipitated using cold acetone and dissolved in 48 μL HENS buffer (250 mM Hepes, pH 7.7, 4 mM EDTA, 1 mM neocuproine, 1% (w/v) SDS), followed by addition of 6 μL 20 mM sodium ascorbate and 6 μL 4 mM biotin-HPDP. This reaction was incubated at room temperature for 1 h. All of the above steps were carried out in a darkroom. Aliquots of each sample was analyzed by SDS-PAGE without boiling, followed by immunoblotting using an anti-biotin antibody (Cell Signaling Technology, Cat # 7075). Then the blot was stripped using strip buffer (15 g/L glycine, 1% SDS, 1% Tween-20, pH 2.2) at room temperature for 30 min, and re-incubated using an anti-His (Santa Cruz Biotechnology, Cat # SC-8036) or anti-GST antibody (Santa Cruz Biotechnology, Cat # SC-138) to verify the input.

### *In vivo S*-nitrosylation Assay

*In vivo* S-nitrosylation assay was performed according previously report^18^. Total cellular proteins were prepared by grinding 7d-old seedlings in liquid nitrogen and then extracted in cold HEN buffer (250 mM HEPES, pH 7.7, 1 mM EDTA, 0.1 mM neocuproine and 1% (v/v) protease inhibitors cocktail). After centrifugation at 13,000 rpm at 4ºC for 20 min, the supernatant was collected and incubated with blocking buffer (250 mM Hepes, pH 7.7, 4 mM EDTA, 1 mM neocuproine, 2.5% (w/v) SDS and 200 mM S-methylmethane thiosulfonate) at 50 ºC for 1 h with frequently vortex. Then the samples were precipitated with cold acetone, followed by washing with acetone for three times, and then dissolved in 240 μL of HEN buffer supplemented with 1% (w/v) SDS. After adding 20 μL of 500 mM sodium ascorbate and 20 μL of 4 mM biotin-HPDP, the samples were incubated at room temperature for 1 h, followed by precipitation and washing with cold acetone. Then the pellet was dissolved in 300 μL 250 mM HEPES buffer and neutralized with 800 μL of neutralization buffer (25 mM HEPES, pH 7.7, 100 mM NaCl, 1 mM EDTA, and 0.5% Triton X-100), followed by adding 30 μL streptavidin beads (Thermo Scientific, Cat #: 29202) and incubated at 4 ºC for 16 h. After washing the beads three times with the buffer (25 mM HEPES, pH 7.7, 600 mM NaCl, 1 mM EDTA, and 0.5% Triton X-100), the samples were then subjected to SDS-polyacrylamide gel electrophoresis and analyzed by immunoblotting.

### DAN Assay

The S-nitrosylated cysteine residue numbers was measured by the 2,3-diaminonaphthalene (DAN) assay, described previously^33^. In brief, around 200 μg recombinant His-IAA17 protein was treated with 200 μM GSNO at room temperature in the darkroom for 1 h. The samples were washed twice with 1 mL PBS to remove excess GSNO, followed by adding 300 μL of 200 mM HgCl_2_ and 200 mM DAN for 30 min in the darkroom, allowing to converse DAN into fluorescent 2, 3-naphthyltriazole (NAT). In order to stop the reaction, 15 μL of 2.8 M NaOH was adding and incubated for 5 min at room temperature. Then the fluorospectrometer (Nano Drop 3300, Thermo Scientific) was used to measure the excitation spectrum of UV (365 ± 10 nm) and emission wavelength (450 nm), that emitted from the NAT fluorescent signal. After the detect, the relative DAN-NAT conversion rate was calculated as the fluorescent intensity of NAT per μM protein.

### Mass Spectrometry

Identification of S-nitrosylated cysteine residues was performed by mass spectrometric using the SNOSID method with minor modifications^18^. Briefly, approximately 100 μg His-IAA17 recombinant proteins were treated with 200 μM GSNO and then labeled with biotin-maleimide (Sigma-Aldrich, Cat #: B1267). After biotinylated proteins were precipitated with cold acetone, the proteins were incubated with 400 μL dissolved buffer (20 mM Tris-HCl, pH 7.7, 1 mM EDTA, 0.4% Triton X-100) and then digested by trypsin (Promega, Cat # V5280) at a final concentration of 10 μg/mL at 37 ºC for 16 h. Then PMSF was adding to terminate the reaction, and the sample were incubated with 50 μL streptavidin-agarose beads (Thermo Scientific, Cat # 29202) at room temperature for 2 h. After that, the streptavidin beads were washed three times in 1 mL of washing buffer (20 mM Tris-HCl, pH 7.7, 1 mM EDTA, 0.4% Triton X-100, 600 mM NaCl), followed by washing three times with 1 mL buffer (5 mM ammonium bicarbonate and 20% acetonitrile). The streptavidin-bound peptides were eluted by adding 100 μL of 100 mM β-mercaptoethanol and then concentrated by a SpeedVac vacuum concentrators. The supernatants were concentrated to a final volume 20-30 μL, and then stored at –20 ºC for subsequent analysis.

The trypsin-digested peptides were analyzed by LC-MS/MS using a BioBasic-18 column (0.18×150 mm^2^, Surveyor System, Thermo Inc.) coupled on-line with the ion trap mass spectrometer (LCQ Deca Xp Plus, Thermo Finnigan). The instrument was run in the data-dependent mode with cycling between one full MS scan from *m/z* 400-2000 and MS/MS scans of five most abundant ions using dynamic exclusion. The normalized collision energy was set to 35% for precursor ions fragmentation. Analysis of MS/MS spectra for peptide identification was performed by protein database searching engine with pFIND software as described before^34^. The His-IAA17 sequence was used as a database for searching. The key searching parameters were set as following: (1) a precursor mass tolerance of 2.0 Da; (2) a fragment mass tolerance of 0.5 Da; (3) two mis-cleavages were allowed in the search; (4) Variable modifications: + 15.995 amu on methionine for oxidation, + 451.540 amu on cysteine for biotinylation; and (5) show spectra or peptides when False Discovery Rate (FDR) ≤ 1%. Choose 2*Y/(X+Y), where X indicates the number of hits to target sequences, and Y indicates the number of hits to decoy sequences, respectively.

### Co-immunoprecipitation assay

The Co-immunoprecipitation (Co-IP) experiments were performed according to previously described methods with minor modifications^32^. Briefly, approximately 20 μg His-IAA17 recombinant protein was incubated with 200 μM GSNO at room temperature in the dark condition for 2 h, then the proteins were washed twice with 1 mL PBS to remove excess GSNO. The 7d-old seedlings of *TIR1-Myc* transgenic plants were grinded in liquid nitrogen, and then extracted in lysis buffer (50 mM Tris-HCl, pH 7.5, 150 mM NaCl, 10 mM MgCl_2_, 10% (v/v) glycerol, 0.1% (v/v) Nonidet P-40, 1 mM phenylmethylsulfonyl fluoride, 1% (v/v) protease inhibitors cocktail (Sigma-Aldrich, P9599). The extracts were cleared by centrifugation at 14,000g for 15 min. After that, the supernatants extracts containing 1.0–2.0 mg total cellular proteins of TIR1-Myc were incubated with 10 μL anti-Myc antibodies for 1 h at 4 °C with gentle shaking. Then the Dynabeads Protein G (50 μL, ThermoFisher) were added and mixed with His-IAA17 protein, which incubated with or without GSNO as indicated before, meaning while adding the 10 μM IAA as indicated. The reaction was run at 4 °C for 2–3 h with gentle shaking. After that, the beads were washed three times with 1 ml washing buffer (50 mM Tris-HCl, pH 7.5, 150 mM NaCl, 10 mM MgCl_2_, 0.1% (v/v) Nonidet P-40) and then subsequently used for immunoblot.

### *In vitro* turnover assay

The analysis of His-IAA17 protein degradation *in vitro* was performed as described methods with minor modifications^32^. In brief, approximately 10 μg His-IAA17 recombinant protein was incubated with 200 μM GSNO at room temperature in the dark condition for 2 h, then the proteins were washed twice with 1 mL PBS to remove excess GSNO. Total protein extracts were prepared from 7d-old *TIR1-Myc* plants grown in 1/2 MS medium using ice-cold extraction buffer (50 mM Tris-HCl, pH 7.5, 150 mM NaCl, 0.01% (v/v) Triton X-100 and 1 mM phenylmethanesulfonyl fluoride, 1% (v/v) protease inhibitors cocktail (Sigma-Aldrich, P9599). The extracts were cleared by centrifugation at 14,000g for 15 min. After that, the supernatants extracts containing (1 mg proteins) were mixed with His-IAA17 protein, which incubated with GSNO as indicated before or 50μM MG132 in a total volume of 600 μL. The mixture was incubated at 4 °C with gentle agitation and 100 μL of each sample was collected at the indicated time points and then analyzed by immunoblotting.

### Quantitative reverse transcription-PCR (qRT-PCR)

Total RNA was prepared using the RNeasy Plant Mini Kit (Qiagen). Quantitative reverse transcription-PCR (qRT-PCR) was performed using the iTaqTM Universal SYBR® Green Supermix (Bio-Rad) according to the manufacturer’s instructions. The reactions were run in a CFX96 REAL-Time PCR Detection System (Bio-Rad). The relative expression level of the target genes was analyzed with the delta-delta Ct method and normalized to the expression level of ACT7. All of the experiments were repeated for at least twice (two biological repeats with three technical repeats for each experiment). The primers used for qRT-PCR are listed in Supplementary Table 1.

## REFERENCE

1 Morffy, N. & Strader, L. C. Structural Aspects of Auxin Signaling. Cold Spring Harb. Perspect. Biol. 14,(2022).

2 Jing, H. & Strader, L. C. Structural biology of auxin signal transduction. In Plant Structural Biology: Hormonal Regulations. J. Hejatko, ed., 49-66,(2018).

3 Figueiredo, M. R. A. & Strader, L. C. Intrinsic and extrinsic regulators of Aux/IAA protein degradation dynamics. Trends Biochem. Sci. 47, 865–874,(2022).

4 Powers, S. K. & Strader, L. C. Regulation of auxin transcriptional responses. Dev. Dyn. 249, 483–495,(2020).

5 Gupta, K. J. et al. Regulating the regulator: nitric oxide control of post-translational modifications. New Phytol. 227, 1319–1325,(2020).

6 Feng, J., Chen, L. & Zuo, J. Protein S-Nitrosylation in plants: Current progresses and challenges. J Integr Plant Biol 61, 1206–1223,(2019).

7 Gupta, K. J. et al. Nitric oxide regulation of plant metabolism. Mol. Plant 15, 228–242,(2022).

8 Kolbert, Z. et al. A forty year journey: The generation and roles of NO in plants. Nitric Oxide-Biol Ch 93, 53–70,(2019).

9 Fernando, V. et al. S-Nitrosylation: An Emerging Paradigm of Redox Signaling. Antioxidants-Basel 8,(2019).

10 Pande, A. et al. Phytohormonal Regulation Through Protein S-Nitrosylation Under Stress. Front. Plant. Sci. 13,(2022).

11 Zhang, J. et al. Recent Progress in Protein S-Nitrosylation in Phytohormone Signaling. Plant Cell Physiol. 60, 494–502,(2019).

12 Fernandez-Marcos, M., Sanz, L., Lewis, D. R., Muday, G. K. & Lorenzo, O. Nitric oxide causes root apical meristem defects and growth inhibition while reducing PIN-FORMED 1 (PIN1)-dependent acropetal auxin transport. Proc. Natl. Acad. Sci. USA 108, 18506–18511,(2011).

13 Terrile, M. C. et al. Nitric oxide influences auxin signaling through S-nitrosylation of the Arabidopsis TRANSPORT INHIBITOR RESPONSE 1 auxin receptor. Plant J. 70, 492–500,(2012).

14 Shi, Y. F. et al. Loss of GSNOR1 Function Leads to Compromised Auxin Signaling and Polar Auxin Transport. Mol. Plant 8, 1350–1365,(2015).

15 Iglesiasa, M. J. et al. Regulation of SCFTIR1/AFBs E3 ligase assembly by S-nitrosylation of Arabidopsis SKP1-like1 impacts on auxin signaling. Redox Biol 18, 200–210,(2018).

16 Gray, W. M., Kepinski, S., Rouse, D., Leyser, O. & Estelle, M. Auxin regulates SCF^TIR1^-dependent degradation of AUX/IAA proteins. Nature 414, 271–276,(2001).

17 Feng, J. et al. S-nitrosylation of phosphotransfer proteins represses cytokinin signaling. Nat Commun 4, 1529,(2013).

18 Zhan, N. et al. S-Nitrosylation Targets GSNO Reductase for Selective Autophagy during Hypoxia Responses in Plants. Mol. Cell 71, 142–154 e146,(2018).

19 Pagnussat, G. C., Simontacchi, M., Puntarulo, S. & Lamattina, L. Nitric oxide is required for root organogenesis. Plant Physiol. 129, 954–956,(2002).

20 Pagnussat, G. C., Lanteri, M. L. & Lamattina, L. Nitric oxide and cyclic GMP are messengers in the indole acetic acid-induced adventitious rooting process. Plant Physiol. 132, 1241–1248,(2003).

21 Otvos, K. et al. Nitric oxide is required for, and promotes auxin-mediated activation of, cell division and embryogenic cell formation but does not influence cell cycle progression in alfalfa cell cultures. Plant J. 43, 849–860,(2005).

22 Ni, M. et al. Excessive Cellular S-nitrosothiol Impairs Endocytosis of Auxin Efflux Transporter PIN2. Front. Plant. Sci. 8,(2017).

23 Lozano-Juste, J. & Leon, J. Nitric Oxide Regulates DELLA Content and PIF Expression to Promote Photomorphogenesis in Arabidopsis. Plant Physiol. 156, 1410–1423,(2011).

24 Chen, L. et al. Nitric oxide negatively regulates gibberellin signaling to coordinate growth and salt tolerance in Arabidopsis. J Genet Genomics 49, 756–765,(2022).

25 Liu, W. Z. et al. Cytokinins can act as suppressors of nitric oxide in Arabidopsis. Proc. Natl. Acad. Sci. USA 110, 1548–1553,(2013).

26 Albertos, P. et al. S-nitrosylation triggers ABI5 degradation to promote seed germination and seedling growth. Nat. Commun. 6,(2015).

27 Ayyar, P. V. Uncovering the Role of S-Nitrosylation in Jasmonic Acid Signalling During the Plant Immune Response. Ph.D. thesis. Edinburgh: IMPS, The University of Edinburgh.,(2016).

28 Gray, W. M. et al. Identification of an SCF ubiquitin-ligase complex required for auxin response in Arabidopsis thaliana. Genes Dev. 13, 1678–1691,(1999).

29 Chen, L. et al. Transnitrosylation Mediated by the Non-canonical Catalase ROG1 Regulates Nitric Oxide Signaling in Plants. Dev. Cell 53, 444–457 e445,(2020).

30 Bechtold, N. & Pelletier, G. In planta Agrobacterium-mediated transformation of adult Arabidopsis thaliana plants by vacuum infiltration. Methods Mol Biol 82, 259–266,(1998).

31 Jing, H. et al. Peptidyl-prolyl isomerization targets rice Aux/IAAs for proteasomal degradation during auxin signalling. Nat Commun 6, 7395,(2015).

32 Jing, H. et al. Regulation of AUXIN RESPONSE FACTOR condensation and nucleo-cytoplasmic partitioning. Nat Commun 13, 4015,(2022).

33 Yang, H. et al. S-nitrosylation positively regulates ascorbate peroxidase activity during plant stress responses. Plant Physiol. 167, 1604–1615,(2015).

34 Li, D. Q. et al. pFind: a novel database-searching software system for automated peptide and protein identification via tandem mass spectrometry. Bioinformatics 21, 3049–3050,(2005).

