## Supplemental data will be used for the link to the file on the manuscript for "*S*-nitrosylation of Aux/IAA protein represses auxin signaling"

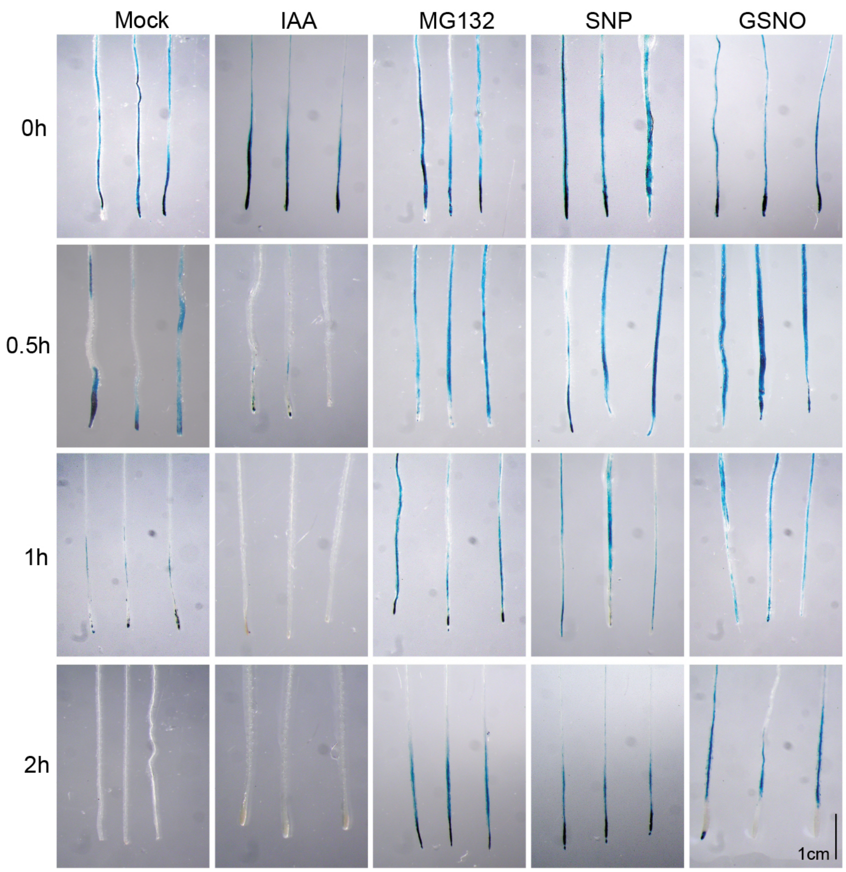


**Supplementary Figure 1. Nitric oxide (NO) regulates HS:AXR3NT-GUS stability.** GUS histochemical staining of 7d-old seedlings containing the HS:AXR3NT-GUS/Col-0 reporter. After heat shock at 37 ºC for 2 h, the seedlings were transferred to media in absence (Mock) or presence of 10 μM IAA, 50 μM MG132, 200 μM SNP, and 200 μM GSNO, incubating for 0-2 h at 22 ºC before GUS staining. Scale bar = 1cm.

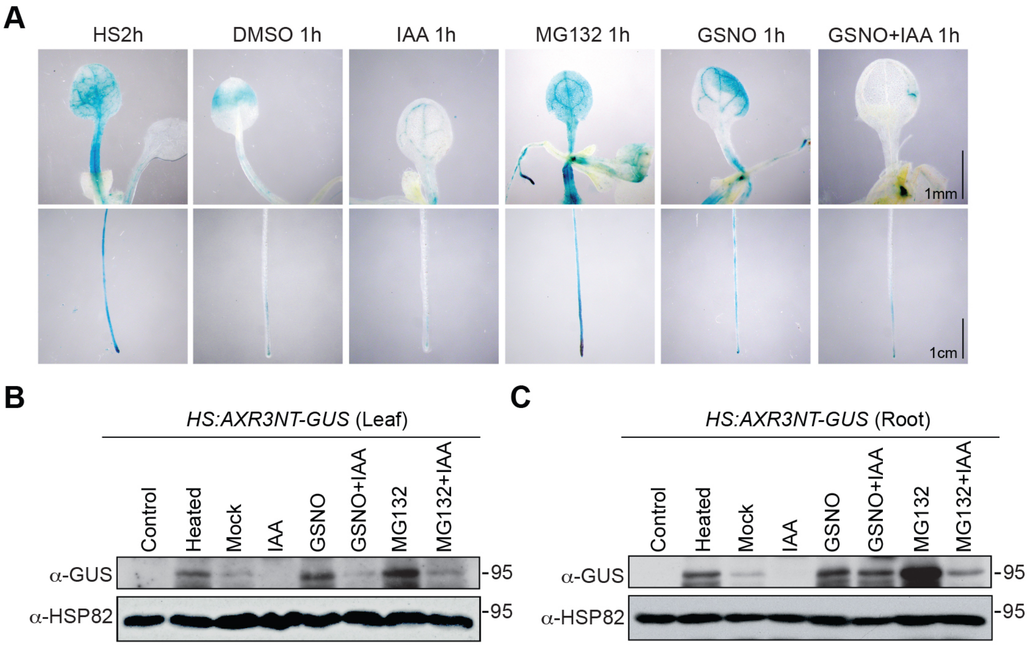


**Supplementary Figure 2. Nitric oxide (NO) inhibits HS:AXR3NT-GUS protein degradation.** (**A**) GUS histochemical staining of 7d-old seedlings leaf and root containing the HS:AXR3NT-GUS/Col-0 reporter. After heat shock at 37 ºC for 2 h, the seedlings were transferred to media in absence (Mock) or presence of 10 μM IAA, 50 μM MG132, 200 μM GSNO, and 200 μM GSNO with 10 μM IAA, incubating for 1 h at 22 ºC before GUS staining. Scale bar = 1mm (leaf), 1cm (root). (**B, C**) Immunoblot analysis of GUS levels from 7d-old HS:AXR3NT-GUS/Col-0 seedlings treated with IAA, GSNO, GSNO with IAA, MG132, and MG132 with IAA, with the indicated concentrations after the end of heat shocked at 37 ºC for 2 h. Anti-HSP82 used for loading control.

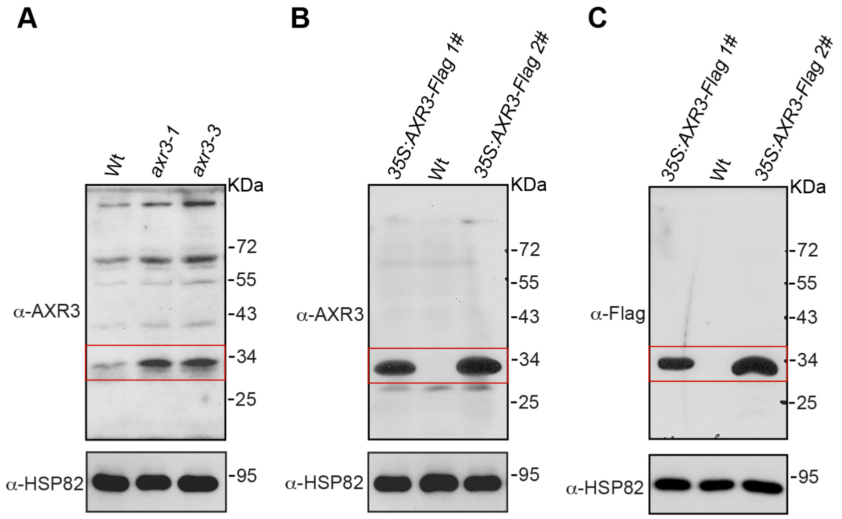


**Supplementary Figure 3. Analysis of anti-IAA17 antibodies.** (**A**) Analysis of the anti-IAA17 antibody in wild-type (Col-0), *axr3-1*, and *axr3-3* mutant plants. A specific band of approximate molecular mass of 34 KDa (indicated by a red box) is detected, which accumulates at a higher level in the *IAA17* gain of function mutant *axr3-1* and *axr3-3.* (**B**) Analysis of anti-IAA17 antibody in wild-type (Col-0) and over expression transgenic plants of *35S:AXR3-Flag*. A band of approximate molecular mass of 34 KDa specific to IAA17 (indicated by a red box) is enriched in *35S:AXR3-Flag* transgenic lines. (**C**) Analysis of anti-IAA17 antibody in wild-type (Col-0) and *35S:AXR3-Flag* transgenic lines using the anti-Flag antibody. This band is also significantly enriched in *35S:AXR3-Flag* transgenic lines, which showing similar results as in the (B).

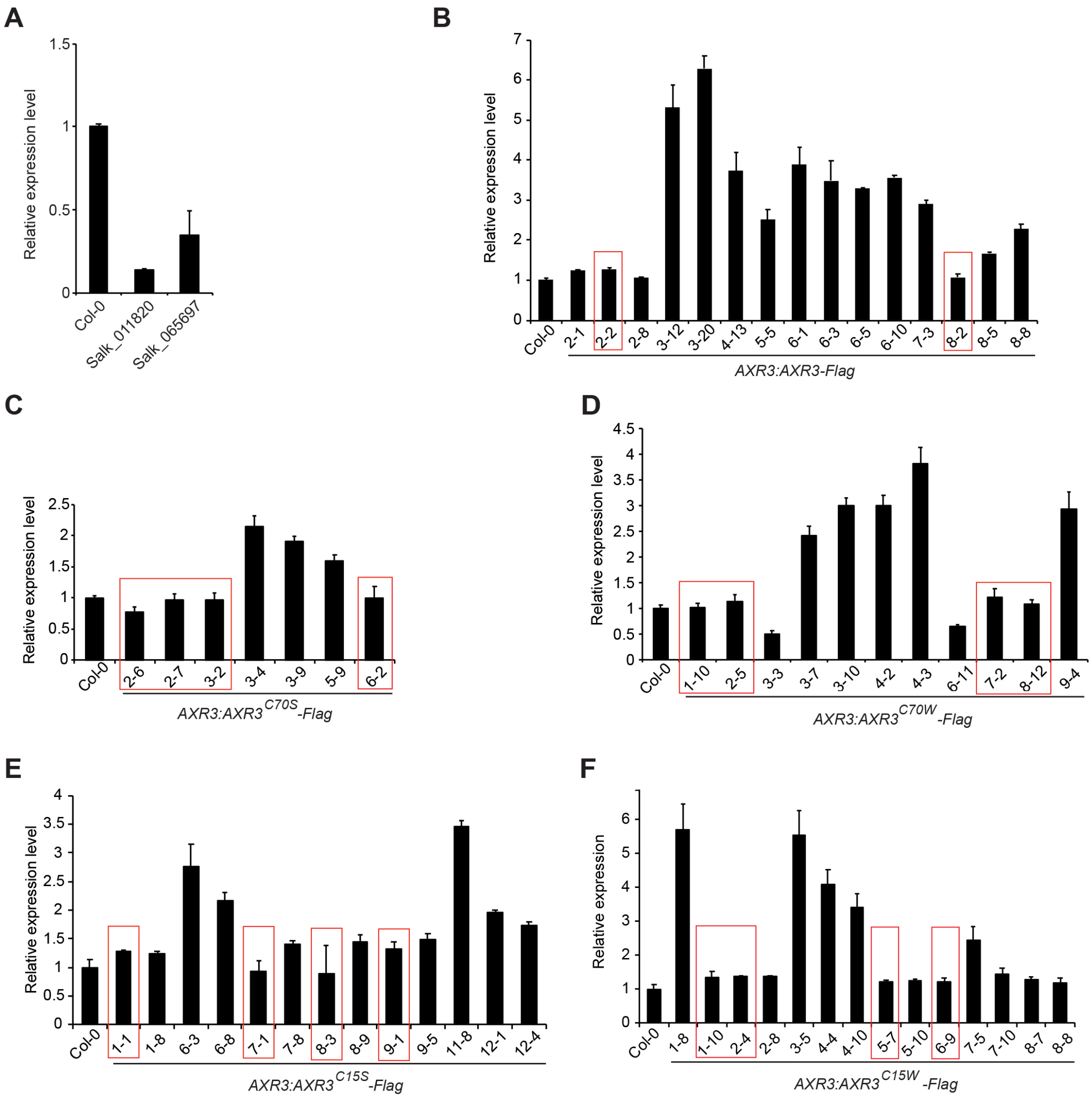


**Supplementary Figure 4. Analysis of the expression levels of *IAA17* genes.** (**A**) Mean relative accumulation of *IAA17* transcript levels in Col-0 and T-DNA insertion lines (Salk_011820 and Salk_065697) as assessed by qRT-PCR. The relative expression level of *IAA17* gene in Col-0 is set at 1.0. Data are mean ± SD from three independent experiments. (**B**) Mean relative accumulation of *IAA17* transcript levels in Col-0 and *AXR3:AXR3-Flag* transgenic lines as assessed by qRT-PCR. The relative expression level of *IAA17* gene in Col-0 is set at 1.0. Data are mean ± SD from three independent experiments. (**C, D**) Mean relative accumulation of *IAA17* transcript levels in Col-0, *AXR3:AXR3$^{C70S}$-Flag, and AXR3:AXR3$^{C70W}$-Flag* transgenic lines as assessed by qRT-PCR. The relative expression level of *IAA17* gene in Col-0 is set at 1.0. Data are mean ± SD from three independent experiments. (**E, F**) Mean relative accumulation of *IAA17* transcript levels in Col-0, *AXR3:AXR3*$^{C15S}$*-Flag, and AXR3:AXR3*$^{C15W}$*-Flag* transgenic lines as assessed by qRT-PCR. The relative expression level of *IAA17* gene in Col-0 is set at 1.0. Data are mean ± SD from three independent experiments.

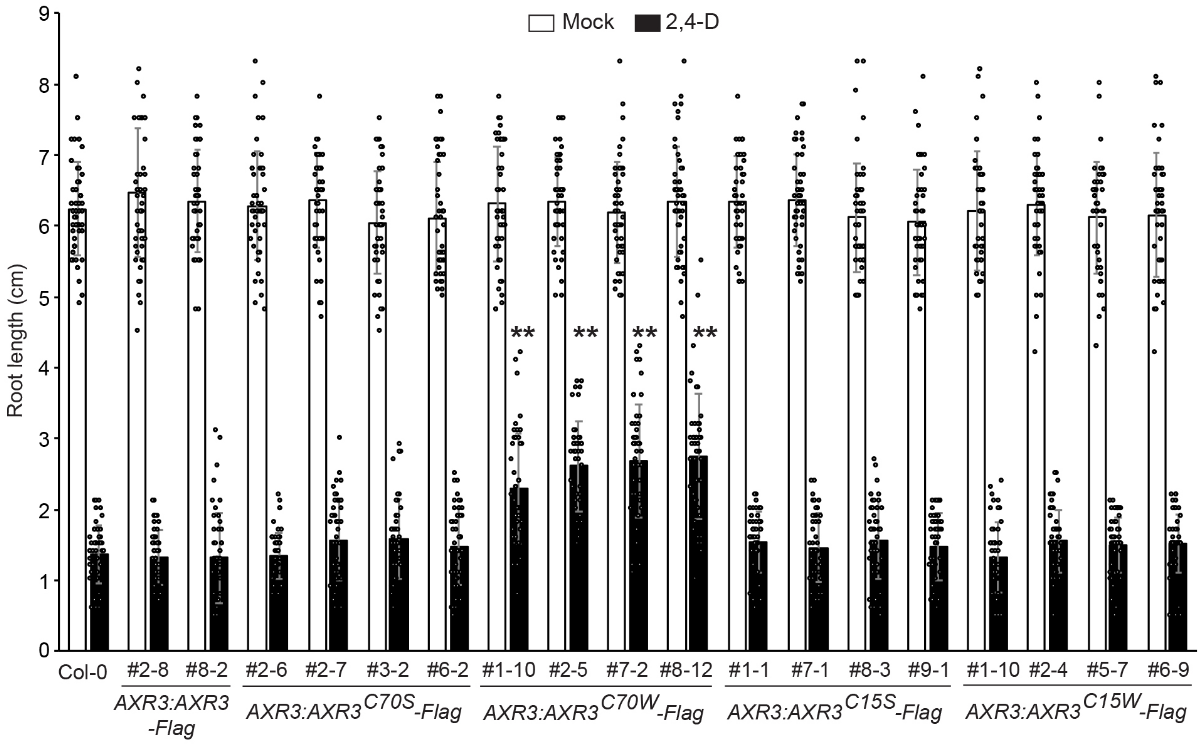


**Supplementary Figure 5. Analysis of the root elongation after 2,4-D treatment.** Mean primary root lengths of 6d-old Col-0, *AXR3:AXR3-Flag*, *AXR3:AXR3$^{C70S}$-Flag*, *AXR3:AXR3$^{C70W}$-Flag, AXR3:AXR3$^{C15S}$-Flag,* and *AXR3:AXR3$^{C15W}$-Flag* seedlings grown on media supplemented with mock (EtOH) or 10 nM 2,4-D after growing 5 days on 1/2 MS media with sucrose. n = 50 biologically independent seedlings were examined in root lengths. Data are mean ± SD from three independent experiments and gray dots represent the individual values. The statistical significance was determined by a two-sided Student's *t*-test (Paired two sample for means). $^{**}P < 0.01$ when compared to Col-0.

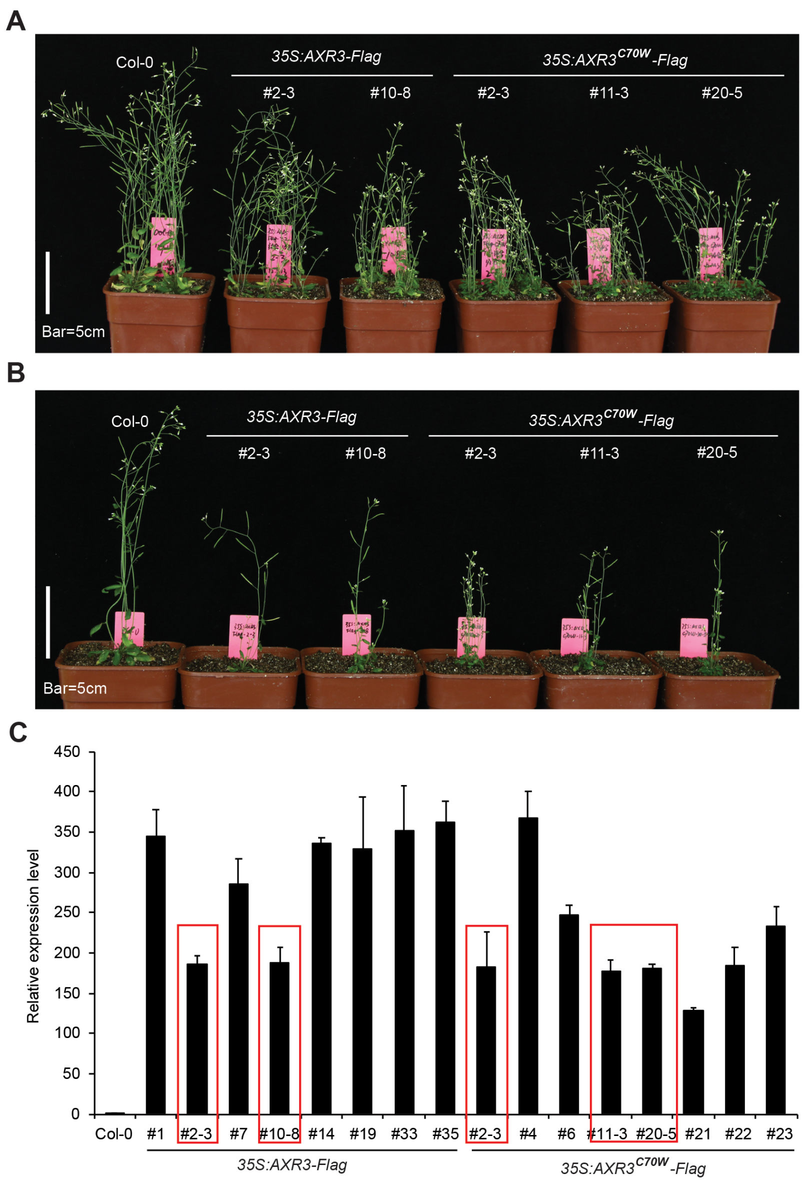


**Supplementary Figure 6. Phenotypes of the overexpression *IAA17*$^{C70W}$ transgenic plants.** (**A, B**) Photograph of 40d-old Col-0, *35S:AXR3-Flag*, and *35S:AXR3*$^{C70W}$*-Flag* plants. Scale bar = 5 cm. (**C**) Mean relative accumulation of *IAA17* transcript levels in Col-0, *35S:AXR3-Flag*, and *35S:AXR3*$^{C70W}$*-Flag* transgenic lines as assessed by qRT-PCR. The transgenic lines showing similar transcript levels indicated as a red box were used to analysis the growth phenotype (A, B). The relative expression level of *IAA17* gene in Col-0 is set at 1.0. Data are mean ± SD from three independent experiments.
